# Structural and functional insight into the plant unique multimodular triphosphosphate tunnel metalloenzymes of *Arabidopsis thaliana*

**DOI:** 10.1101/2022.03.18.484887

**Authors:** Marta Pesquera, Jacobo Martinez, Kai Wang, Manuel Hofmann, Sylvain Loubéry, Priscille Steensma, Michael Hothorn, Teresa B. Fitzpatrick

## Abstract

Triphosphate tunnel metalloenzymes (TTMs) are found in all biological kingdoms and have been characterized in microorganisms and animals. Members of the TTM family already characterized have divergent biological functions and act on a range of triphosphorylated substrates (RNA, thiamine triphosphate, inorganic polyphosphate). TTM proteins in plants have received considerably less attention and are unique in that some homologs harbor additional domains including a P-loop kinase and transmembrane domain. Here we report on structural and functional aspects of the multimodular TTM1 and TTM2 of *Arabidopsis thaliana*. Tissue and cellular microscopy studies show that both AtTTM1 and AtTTM2 are expressed in actively dividing (meristem) tissue and are tail-anchored proteins at the outer mitochondrial membrane - mediated by the single transmembrane domain at the C-terminus, supporting earlier studies. Crystal structures of AtTTM1 in the presence and absence of a non-hydrolyzable ATP analog reveal a catalytically incompetent TTM tunnel domain tightly interacting with the P-loop kinase domain that is locked in an inactive conformation. Structural comparison reveals that a helical hairpin may facilitate movement of the TTM domain thereby activating the kinase. Genetic studies show that At*TTM2* is important for the developmental transition from the vegetative to the reproductive phase in Arabidopsis, whereas its closest paralog At*TTM1* is not. Rational design of mutations based on the 3D structure demonstrates that both the P-loop kinase and TTM tunnel modules of AtTTM2 are required for the developmental switch.

## INTRODUCTION

TRIPHOSPHATE TUNNEL METALLOENZYMEs (TTMs) are found in all biological kingdoms. The family is characterized by an eight-stranded β-barrel tunnel domain (1) that harbors a set of basic amino acids involved in the binding of various triphosphate substrates and one or two conserved glutamic acid residues involved in the coordination of one (2) or two (3) divalent metal ions required for catalysis. M**Error! Bookmark not defined**.embers of this superfamily accept a wide range of substrates comprising nucleotides and organophosphates, including RNA (1), inorganic triphosphate (3-5), thiamine triphosphate (3,6) and cyclic AMP (cAMP) (7,8). While most TTM enzymes hydrolyze triphosphate substrates, the TTM protein Vtc4p catalyzes the synthesis of inorganic polyphosphate from ATP (2). TTM domains may occur as stand-alone enzymes, or fused to for example inorganic polyphosphate binding CHAD domains (3,9,10) or to inositol pyrophosphate sensing SPX domains (11).

In the model plant *Arabidopsis thaliana*, there are three *TTM* genes (At*TTM1-3*) (12). All three are reported to carry the TTM domain characteristic of the family, sometimes referred to as CYTH, defined by the two founding members <u>Cy</u>aB adenylyl cyclase from *Aeromonas hydrophila* and mammalian <u>th</u>iamine triphosphatase (13). AtTTM3 was the first member to be characterized and originally thought of as a potential adenylate cyclase involved in the formation of cyclic AMP, the presence of which in plants is controversial since decades (14), but was later reported to be a tripolyphosphatase involved in root development (3,12). More recently, AtTTM3 has been shown to be part of a bicistronic transcript that harbors an ortholog of a cell cycle regulator (15). In contrast to AtTTM3, AtTTM1 and AtTTM2 form part of the plant unique cluster of TTM proteins that exclusively occur in a tandem arrangement with a N-terminal P-loop nucleoside triphosphate phosphotransferase/kinase domain (4). While structural information is available for the single TTM domain protein AtTTM3 (16), the architectural arrangement of multimodular AtTTM members is unknown. Further, both AtTTM1 and 2 are reported to be tail-anchored to the outer mitochondrial membrane and have been previously characterized as pyrophosphatases (17-19), rather than acting on triphosphorylated substrates as is characteristic of the TTM family. Conditional phenotypes have been reported for loss of these multimodular TTM proteins with AtTTM1 implicated in abscisic acid-induced senescence, while AtTTM2 acts as a negative regulator in salicylic acid mediated defense responses (18-20). Although, it is not known whether these proteins play a role in developmental processes.

Here we report structural and functional analyses of AtTTM1 and AtTTM2. Commonalities among AtTTM1 and AtTTM2 are that they are expressed largely in actively dividing tissue and are tail-anchored to the outer mitochondrial membrane in support of earlier studies. Crystal structures of AtTTM1 reveal an unusual tandem arrangement of the TTM and kinase domains, with the TTM domain being inserted into the active site of the kinase, possibly keeping it in an inactive conformation. Interestingly, AtTTM2, unlike AtTTM1, has a distinct role to play in the transition from the vegetative to the reproductive state (bolting) in Arabidopsis which requires both the TTM and kinase domains to be functional.

## RESULTS

### Cellular and tissue localization of AtTTM1 and AtTTM2 largely overlap

Both AtTTM1 (At1g73980) and AtTTM2 (At1g26190) have previously been reported to be tail-anchored to the outer mitochondrial membrane (18). Here, examination of the pattern of expression of translational fusions of either AtTTM1/2 with YFP at the N terminus in mesophyll protoplasts isolated from Arabidopsis by confocal fluorescence microscopy is also indicative of mitochondrial localization, as there was no overlap with chlorophyll autofluorescence nor a diffuse pattern indicative of localization to the cytosol, as seen for YFP alone (Fig. 1A, B). The mitochondrial localization was confirmed by co-staining with MitoTracker® Red CMXRos (Fig. 1A, B). On the other hand, a diffuse pattern of fluorescence was observed with both AtTTM1-YFP and AtTTM2-YFP indicative of cytosolic localization (Fig. 1A, B), and suggests that mitochondrial localization in AtTTM1 and AtTTM2 is disrupted when YFP is fused to the C-terminus. This corroborates the previous work on AtTTM1 and AtTTM2 on localization to the mitochondria (18). We further examined localization *in planta* by generating stable transgenic Arabidopsis lines. Fluorescence was examined in hypocotyl or leaf epidermal cells as well as root tips of young seedlings and the punctate pattern of YFP-AtTTM1 or YFP-AtTTM2 in these lines was consistent with mitochondrial localization and the pattern overlapped upon co-staining with MitoTracker® Red CMXRos (Fig. 1C). Magnified root images show that YFP fluorescence of YFP-AtTTM1 or YFP-AtTTM2 displays a ring pattern around the stained mitochondria (Fig. 1C, bottom panels), characteristic of outer mitochondrial membrane proteins (21). That the single transmembrane domain of each protein was required for anchoring to the mitochondria was verified by examining the fluorescence pattern of transgenic lines expressing YFP fused to truncated forms of AtTTM1 or AtTTM2 without the transmembrane domain (YFP-AtTTM1ΔTM, YFP-AtTTM2ΔTM, respectively), in which case a diffuse pattern indicative of localization to the cytosol was observed (Fig. 1C). A protease protection assay with thermolysin on intact mitochondria isolated from YFP-AtTTM1 or YFP-AtTTM2 lines indicated that both proteins are exposed on the surface of the mitochondria (Fig. S1). We also examined the tissue expression pattern of both genes using promoter-GUS reporter lines (*p*At*TTM1:GUS*) and (*p*At*TTM2:GUS*). Representative pictures of *GUS* expression show that both At*TTM1* and At*TTM2* are expressed in shoot and root apices and are pronounced in actively dividing tissue—emerging leaves and lateral root primordia (Fig. 1D, E). Notably, At*TTM1* promoter activity appears to be stronger than At*TTM2* (Fig. 1D, E). Furthermore, and in contrast to At*TTM2*, At*TTM1* promoter driven activity was found in vascular tissue and inflorescences (Fig. 1D, E). Collectively, our data support earlier studies in that AtTTM1 and AtTTM2 are anchored to the outer mitochondrial membrane through a transmembrane domain at their respective C-termini and are both localized to actively dividing tissue.

**Figure 1.**
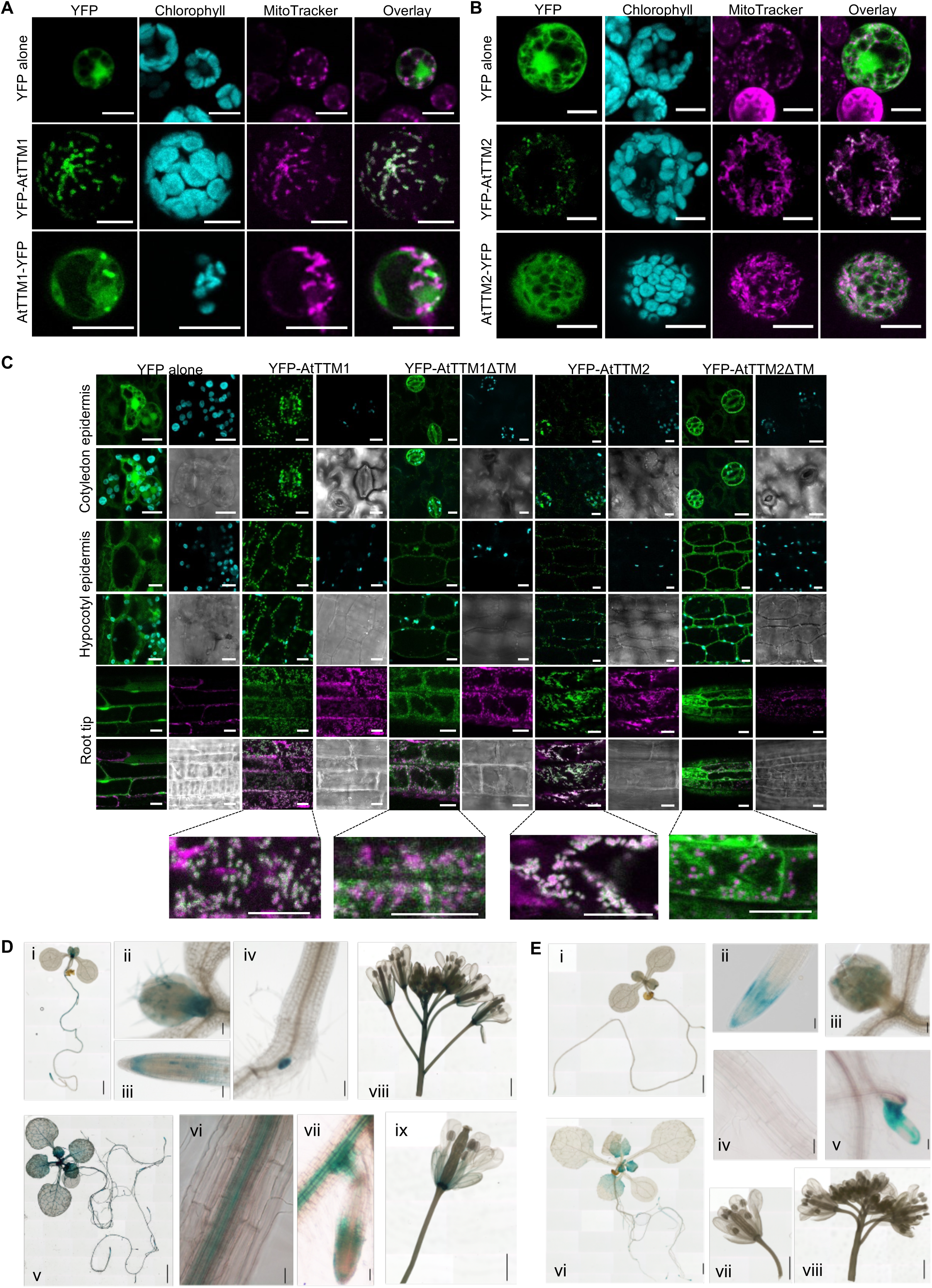
Cellular and tissue expression of AtTTM1 and AtTTM2 in Arabidopsis. A and B, Representative confocal microscope images illustrating transient expression of YFP alone, YFP-AtTTM1, AtTTM1-YFP, YFP-AtTTM2 and AtTTM2-YFP in mesophyll protoplasts. The columns from left to right show YFP (green), chlorophyll (blue) and MitoTracker (magenta) and an overlay of YFP and MitoTracker fluorescence, respectively. Scale bars: 10 µm. C, Confocal images of the epidermis of cotyledons or hypocotyl, and root tip of 3-day-old seedlings expressing YFP alone, YFP-AtTTM1 or YFP-AtTTM2 and a truncated version of YFP-AtTTM1 or YFP-AtTTM2 without the transmembrane domain (YFP-AtTTM1ΔTM and YFP-AtTTM2ΔTM, respectively). The fluorescence coloring is the same as shown in A and B. The four-panel squares from left to right and top to bottom show YFP, chlorophyll, overlay of YFP and chlorophyll fluorescence and bright field in epidermal cells of cotyledons and hypocotyl. The lower panel group show YFP, MitoTracker, overlay of YFP and MitoTracker fluorescence, and bright field in roots. Scale bars: 10 µm. D, Representative images of staining for At*TTM1* promoter driven *GUS* expression. Top panels show a 1-week old whole seedling (i) and the corresponding shoot apical meristem region (ii), the root tip (iii) and differentiation zone showing lateral root emergence (iv). The lower panels are images of a whole 2-week old seedling (v), the corresponding root elongation zone (vi), as well as differentiation zone showing vasculature (vii). The top and bottom right panels are images of an inflorescence (viii) and an open flower (ix), respectively. Scale bar = 1 mm for (i), (v), (viii) and (ix); 100 µm for (ii); 50 µm for (iii), (iv) and (vii); 2.5 µm for (vi). E, Representative images of staining for At*TTM2* promoter driven *GUS* expression of a 1-week old whole seedling (i) and the corresponding root tip (ii), shoot apical meristem region (iii), the elongation zone (iv) and differentiation zone showing lateral root emergence (v). Images of a whole 2-week old seedling (vi), an open flower (vii) and inflorescence (viii). Scale bar = 1 mm for (i), (vi), (vii) and (viii); 100 µm for (iii); 50 µm for (ii) and (v); 2.5 µm for (iv). In all cases, Arabidopsis plants were grown under a 16 hr photoperiod.

### Arabidopsis TTM1 harbors a catalytically inactive TTM domain

We expressed and purified the tandem kinase-TTM domain modules from AtTTM1 (residues 20-412) and AtTTM2 (residues 20-412) for structural analysis. We could only obtain crystals for selenomethionine-labeled AtTTM1. The structure of AtTTM1 was solved to 2.7 Å resolution (Table S1). AtTTM1 folds into a N-terminal kinase domain that is connected to the C-terminal TTM domain with a short, rigid, helical linker (shown in gray in Fig. 2A). The two domains share an extensive interface, with loops from the β-tunnel domain inserting into the putative substrate binding region of the P-loop kinase domain (Fig. 2A, see below). As previously seen for many other TTM proteins, a buffer molecule (citrate) is found bound in the TTM domain center, making extensive contacts with the conserved arginine and lysine residues lining the tunnel walls (Fig. 2A-C) (3,4,12). The kinase domain contains a Walker A motif (P-loop) (22) and Walker B motif (GxxxxGK(S/T and hhhhE), which are predicted to be involved in substrate binding and coordination of a metal ion, respectively (23). In line with this, a phosphate ion is found in proximity to the P-loop motif of the kinase (Fig. 2A). A structural homology search with the program DALI (24) returned different bacterial and archaeal TTM domains as top hits (DALI Z-scores ∼16-13). Among them is the crystal structure of TTM from the archaeon *Sulfolobus acidocaldarius* (SaTTM) (4). Structural superposition of the TTM domain of AtTTM1 with SaTTM revealed that many basic residues lining the tunnel walls are conserved, as is the Tyr404 in the C-terminal plug helix (Fig. 2C, D) (3,4,12). Two conserved glutamate residues in SaTTM from the N-terminal ExExK motif, involved in the coordination of the two metal cofactors, are replaced by glutamines in both AtTTM1 and 2 (Fig. 2D). Consequently, no metal co-factor bound structure of AtTTM1 could be obtained, suggesting the TTM domain of AtTTM1 may not be able to hydrolyze phosphorylated substrates. However, the conserved tunnel domain shape and the presence of conserved basic amino acids along the tunnel walls implicate this domain in the coordination of a (tri)phosphate-containing metabolite.

**Figure 2.**
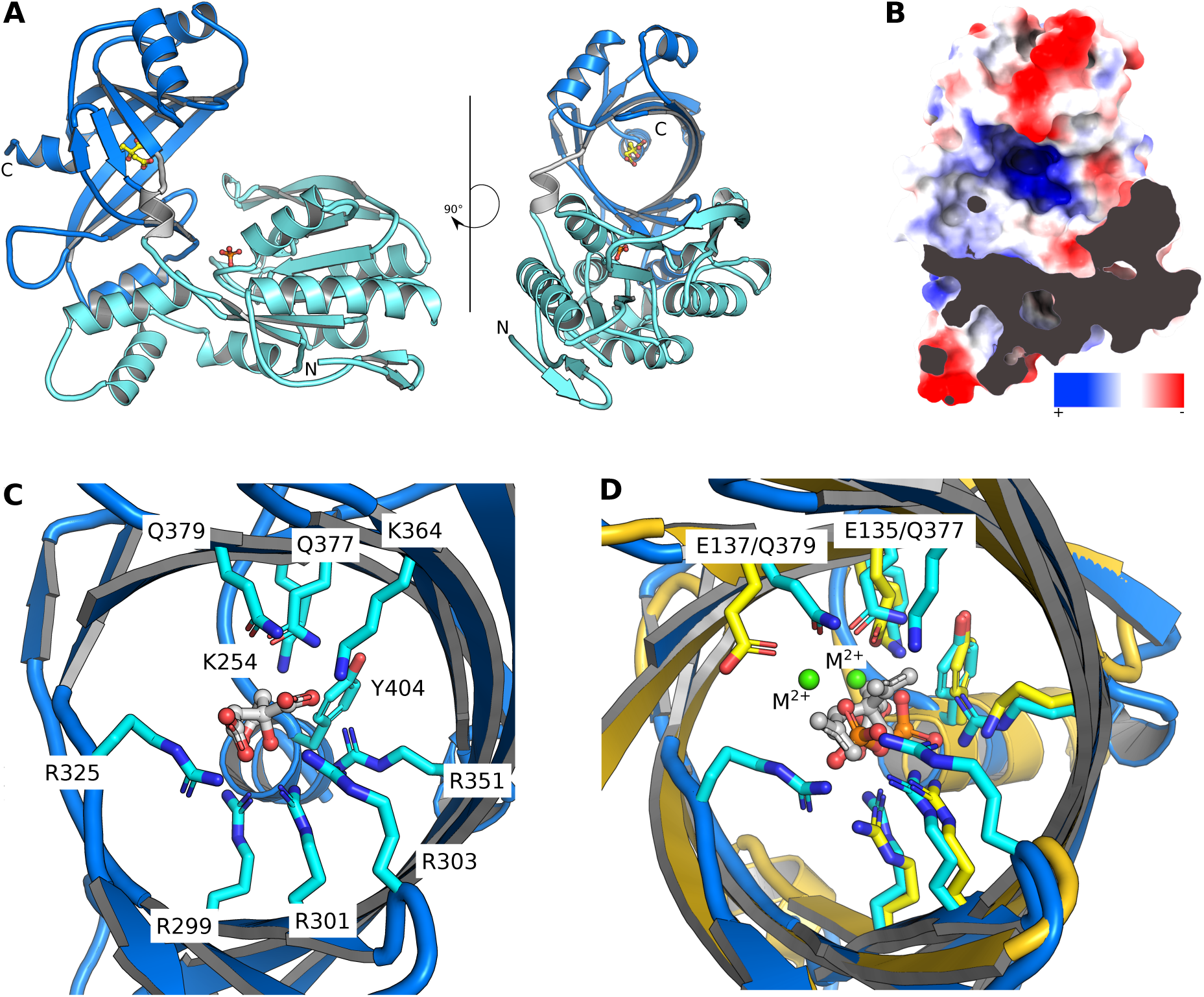
The N-terminal kinase domain of AtTTM1 tightly interacts with a catalytically inactive TTM domain. A, Ribbon diagram of AtTTM1 (residues 20-412) with the N-terminal kinase domain and the C-terminal triphosphate tunnel metalloenzyme (TTM) domain shown in cyan and blue, respectively. An inorganic phosphate ion bound to the P-loop of the kinase domain and a citric acid molecule in the TTM domain tunnel center are highlighted in bond representation (in yellow). B, View of the highly basic tunnel center of the C-terminal TTM domain. An electrostatic potential has been mapped onto the molecular surface of AtTTM1 colored from red (negative charges) to blue (positive charges). The N-terminal kinase domain has been omitted for clarity. C, The pseudo-ligand citric acid (in bond representation, in gray) is coordinated by a set of conserved basic amino acids (in bond representation, in cyan) in the tunnel center of the TTM domain. D, Structural superposition of the AtTTM1 TTM domain (in blue) with the tripolyphosphatase from *Sulfolobus acidocaldarius* (PDB-ID 7NS9, root mean square deviation is ∼2.1 Å comparing 152 corresponding C_*α*_ atoms, in yellow) reveals that Glu135 and Glu137 involved in coordination of two metal ions (in green) required for catalysis of tripolyphosphate (in bond representation, in orange) in SaTTM are replaced by glutamine in AtTTM1.

### The TTM domain may regulate the phosphotransferase activity of the P-loop kinase domain

The presence of a phosphate ion in the kinase domain of AtTTM1 (Fig. 2A) prompted us to co-crystallize the enzyme in the presence of nucleoside triphosphate (NTP) substrates. We obtained a structure in complex with the non-hydrolyzable ATP analog adenosine-5’-[(β,γ)- methyleno]triphosphate (APPCP) to 2.65 Å resolution. We found the triphosphate substrate bound to the NTP binding pocket of the kinase domain (Fig. 3A). The γ-phosphate of the substrate analog is well defined in the electron density and positioned ∼16 Å from the citrate pseudo-substrate bound to the tunnel center of the TTM domain (Fig. 3A). This makes a direct phosphotransfer reaction between the kinase and TTM domain unlikely. The γ-phosphate is however pointing towards a proximal binding pocket with a width of ∼13 Å (Fig. 3A). We inspected this proximal substrate binding site in the AtTTM1 kinase domain and located a partially negatively charged surface area in proximity to the APPCP γ-phosphate (Fig. 3B). Importantly the site contains a hydrophobic pocket, deeply inserting into the kinase fold (Fig. 3B). DALI returned several uridine-cytidine kinase (UCK) structures as top hits for the kinase domain of AtTTM1 (DALI Z-scores ∼14-23), supporting the notion that AtTTM1 may harbor small molecule phosphotransferase activity. The crystal structure of the human UCK (25) closely aligns with the kinase domain of AtTTM1 (Fig. 3C). The substrate analog APPCP in AtTTM1 and the ADP reaction product of HsUCK occupy the same binding site (Fig. 3C, cyan and red, respectively), while cytidine monophosphate in HsUCK maps to the proximal binding surface in AtTTM1 (see above, Fig. 3C).

**Figure 3.**
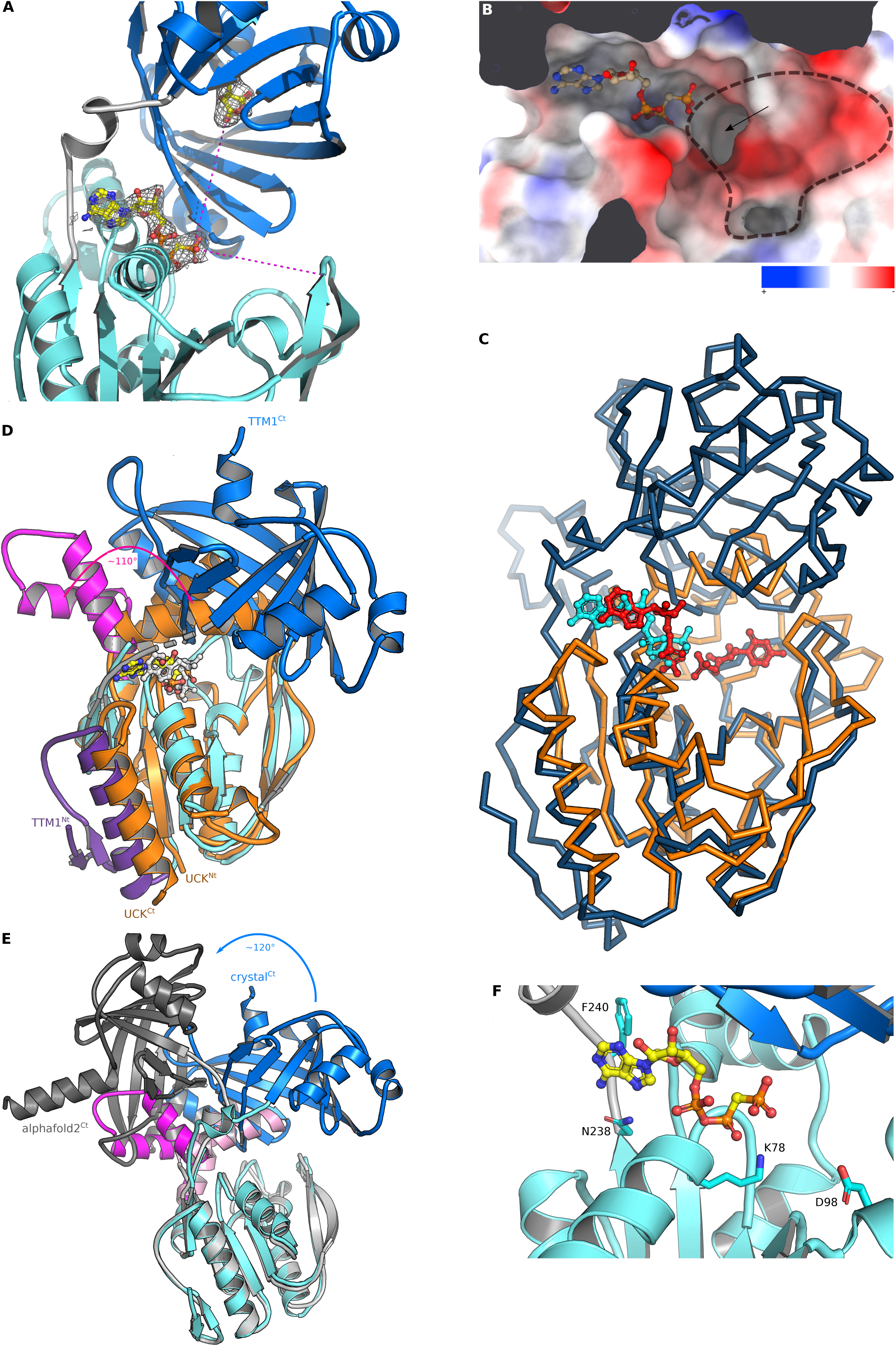
AtTTM1 is a phosphotransferase adopting an auto-inhibited conformation. A, Ribbon diagram (colors as in Fig. 2) of AtTTM1 co-crystallized in the presence of the nucleotide analog adenosine-5’-[(β,γ)- methyleno]triphosphate (APPCP, in bond representation) and including an 2F_o_-2F_c_ omit electron density map contoured at 1.2 s (gray mesh). The distance of the APPCP *γ*-phosphate and the citric acid molecule in the TTM domain is ∼16 Å, the width of the proximal binding site is ∼13 Å (dotted lines, in magenta). B, Surface representation of the substrate binding sites in the AtTTM1 kinase domain. An electrostatic potential has been mapped onto the molecular surface of AtTTM1 colored from red (negative charges) to blue (positive charges). A putative second substrate binding site is highlighted by a dotted line, the hydrophobic pocket discussed in the text is marked by an arrow. C, Structural superposition (root mean square deviation is ∼1.8 Å comparing 120 corresponding C_*?*_ atoms) of AtTTM1 (in dark-blue, C_*?*_-trace) with human UCK (in orange, PDB-ID 1UJ2) reveals the AtTTM1 TTM domain inserted into the uridine/cytidine binding pocket. The APPCP ligand in AtTTM1 and ADP/CMP in HsUCK are shown in cyan and red, respectively (in bond representation). D, Side-view of a structural superposition of the P-loop kinase domains shown in (C). AtTTM1 (blue ribbon diagram) and HsUCK (in orange) closely align, with the exception of a helical hairpin that in HsUCK forms part of the nucleotide binding pocket and that is found 110° rotated outwards in the AtTTM1 crystal structure (AtTTM1 hairpin shown in magenta). Note that the C-terminal helix in HsUCK (in orange) is replaced by an extra N-terminal helix in AtTTM1 (in purple). The nucelotide analogs bound to HsUCK and AtTTM1 kinase domain are shown in bond representation alongside and colored in gray and yellow, respectively. E, Structural superposition of the experimentally determined crystal structure of AtTTM1 (in blue, colors as in (A)) with a theoretical model generated by AlphaFold (https://alphafold.ebi.ac.uk/entry/Q9C9B9) using the kinase domain as reference. The AlphaFold model presents the helical hairpin (in light magenta) flipped inwards, completing the nucleotide binding pocket. The AlphaFold model TTM domain (in gray) appears swung out, away from the second substrate binding site (∼120° rotation). F, Close-up view of the AtTTM1 nucleoside triphosphate binding site bound to APPCP (in yellow). Key interacting residues are shown alongside (in bond representation), the residues corresponding to Lys78 and Asp98 are involved in catalysis in human and bacterial UCKs.

Comparison with the HsUCK structure revealed that the linker connecting the kinase and the tunnel domains of AtTTM1 replaces a conserved C-terminal helix found in HsUCK and other P-loop kinases (Fig. 3D, in gray). Notably, all members of the plant-unique TTM-kinase family harbor an additional N-terminal *α*-helix that replaces the ‘missing’ C-terminal helix and completes the kinase core fold (Fig. 3D, in purple). The AtTTM1 tunnel domain inserts itself into the proximal substrate binding surface (Fig. 3C, D, in blue). This apparently causes a helical hairpin in the kinase domain, normally involved in the coordination of the nucleoside substrate, to be displaced by an ∼ 110° rotation (Fig. 3D, in magenta), thus keeping the kinase domain in an inactive conformation. Comparison with a theoretical structural model of AtTTM1 generated with AlphaFold (https://alphafold.ebi.ac.uk/entry/Q9C9B9) revealed that movement of the TTM domain may enable the helical hairpin to flip towards the active site, thus activating the kinase (Fig. 3E, in magenta). Together, plant TTM-UCK proteins share structural features with known UCK and other P-loop kinases but have tightly integrated an unusual tunnel domain into their active site (26).

Both AtTTM1 and AtTTM2 were initially annotated as UCK proteins (23) but have since been reported to act as pyrophosphatases with inorganic pyrophosphate as the substrate (18). Our structural analyses prompted us to assess if AtTTM1 and 2 harbor phosphotransferase activity. Incubation of AtTTM1 with uridine as substrate in the presence of ATP led to the production of uridine monophosphate (UMP), clearly suggesting that AtTTM1 displays phosphotransferase activity (Fig. S2). A mutant form of AtTTM1 in which lysine 78 involved in coordination of the substrate γ- phosphate and the general base aspartate 98 (25) (Fig. 3F) were replaced by alanine had no detectable activity (Fig. S2). We could detect no phosphotransferase activity for AtTTM2 suggesting that either it does not use uridine as a substrate, in contrast to AtTTM1, or it is not active as isolated and assayed in this manner (Fig. S2).

### AtTTM2 alone is required for the timely transition to the reproductive state in Arabidopsis

We next performed phenotypic analysis of T-DNA insertion lines of At*TTM1* and At*TTM2* under standard growth conditions. For At*TTM1*, PCR analysis established insertion of the T-DNA in exon 3 in SALK_079237, exon 9 in GK_672E02 and intron 10 in SALK_126667 at 551 bp, 2183 bp and 2523 bp, downstream of the translational start codon, respectively. In a previous independent study (18), SALK_079237 is annotated as *ttm1-1* and GK_672E02 as *ttm1-2*, accordingly we annotated SALK_126667 as *ttm1-3* (Fig. 4A). Analysis of the transcript levels of At*TTM1* expression by quantitative real-time RT-PCR (RT-qPCR) confirmed reduced expression in two (*ttm1-1* and *ttm1-3*) out of the three lines (Fig. 4B). In our hands, no incongruent morphological phenotype was observed over the life cycle with any of these lines compared to wild type when grown on soil under standard conditions. In the case of At*TTM2*, PCR analysis established insertion of the T-DNA in exon 3 for SALK_084280 and SALK_145897, and intron 5 for SALK_114669 at 465 bp, 548 bp, and 1159 bp downstream of the translational start codon, respectively. Previously, SALK_145897 was annotated as *ttm2-1* and SALK_114669 as *ttm2-2* (19), accordingly we annotated SALK_084280 as *ttm2-3* (Fig. 4A). RT-qPCR confirmed reduced transcript levels of At*TTM2* in all three lines (Fig. 4B). When grown on soil under standard conditions using a 16 hr photoperiod, these mutant lines showed no incongruent phenotype during the vegetative stage. However, we observed a delay in the transition to the reproductive stage (i.e. bolting) concomitant with an increase in total leaf number in all three *ttm2* mutant lines compared to wild type (Fig. 4C and D). When *ttm1-1* was crossed with *ttm2-2* there was no significant difference in the bolting delay of the *ttm1-1 ttm2-2* double mutants compared to *ttm2-2* alone (Fig. 4C and D). Transformation of *ttm2* with At*TTM2* under the control of its upstream region (*pTTM2:TTM2*) and isolation to homozygosity, restored the timing of the developmental switch (Fig. 4B-D), confirming its involvement in this process. Notably, bolting time was delayed in *ttm2* mutant lines in both 16 hr and 8 hr photoperiods (Fig. S3A). Further, bioactive gibberellin (GA_3_) had no effect on bolting time in *ttm2* compared to wild type (Fig. S3B). The clear morphological phenotype of *ttm2* lines (in contrast to *ttm1*) permitted us to assess *in vivo* if both the kinase and TTM domains are required for functionality. We transformed *ttm2-1* with AtTTM2 isoforms harboring the same mutations as used for the biochemical assays (annotated as *pTTM2*:TTM2 K78A/D98A and *pTTM2*:TTM2 R299A/R301A, respectively). Even though, the mutated AtTTM2 isoforms were expressed at levels similar to (or higher than) the wild type isoform *pTTM2:TTM2* (Fig. S4A), they could not restore bolting or leaf number of *ttm2-1* to that of wild type (Fig. S4B-D). This implies that both the kinase and TTM domain are required for AtTTM2 functionality in Arabidopsis.

**Figure 4.**
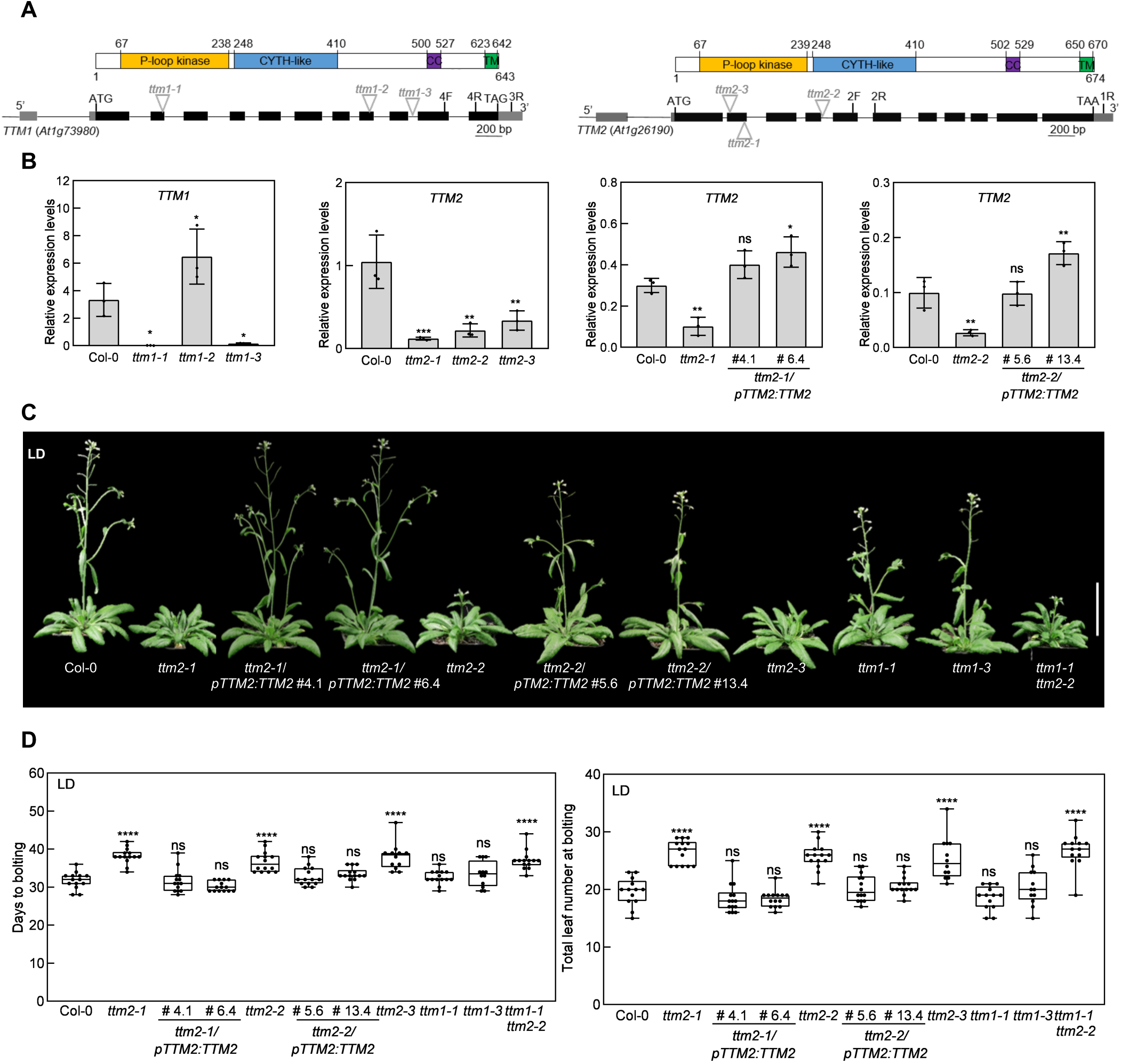
At*TTM2* plays a role in the switch from vegetative to reproductive growth. A, Schematic representation of the gene models of At*TTM1* and At*TTM2* with untranslated regions depicted in gray, exons in black and introns as lines and the localization of the T-DNA insertions in mutant alleles (bp, base-pairs), above which are the corresponding AtTTM1 and AtTTM2 protein predicted domains (CC, coiled-coil; TM, transmembrane domain). B, Quantitative analysis of At*TTM1* or At*TTM2* expression in *ttm1* or *ttm2* mutants and At*TTM2*-complemented lines (*pTTM2:TTM2*) versus wild type (Col-0) relative to *PP2AA3* or *UBC21*. The data are the average of three independent biological replicates. The position of primers used is indicated by 1-4F/R. Error bars represent standard deviation. Statistical relevance was calculated by one-way ANOVA followed by Dunnett’s multiple comparisons test. P<0.05*, P<0.01** and P<0.001***, respectively. C, Representative photographs of the transition to the reproductive stage of the lines as indicated under a 16 hr photoperiod (long day, LD), the scale bar=5 cm. D, Quantification of the switch to the reproductive stage represented by days to bolting and total leaf number (rosette and cauline leaves). Box-plot graphs show the median (horizontal bar in the box) and the minimum and maximum of the data (whisker extremities). One representative replicate of three biological replicates is shown (n = 12-14 per replicate). Statistical relevance was calculated by one-way ANOVA followed by Dunnett’s multiple comparisons test, P > 0.05 ns, P<0.0001****. In all cases ns refers to no significance.

## DISCUSSION

Here a combination of cellular and molecular experiments demonstrates that the multimodular TTM proteins of Arabidopsis overlap considerably in relation to tissue and subcellular localization. Crystallographic analysis of AtTTM1, a founding member of the plant-unique family of multimodular TTM proteins (4) revealed an unusual domain architecture. The N-terminal kinase domain shows strong structural similarity with eukaryotic and prokaryotic UCK’s, is able to bind a nucleoside triphosphate substrate and can catalyze phosphotransfer reactions *in vitro*. The TTM domain however protrudes into the proximal binding site of the kinase domain that normally harbors the cytidine/uridine substrate. AtTTM1 and 2 may thus either bind substrates other than cytidine/uridine or may represent bona fide UCK enzymes in an auto-inhibited state. Indeed, ligand-binding to the C-terminal TTM domain, post-translational modification of AtTTM1/2 or protein-protein interactions, may lead to a structural rearrangement of the kinase domain. Recent studies *in vitro* demonstrated phosphorylation of AtTTM1 by a suite of recombinant mitogen activated protein (MAP) kinases and phosphorylation was shown to mediate AtTTM1 functionality (20). Future studies will establish if this facilitates the conformational change in the protein. As illustrated in Fig. 3E, the TTM domain may move away from the substrate binding site, enabling the *α*-helical hairpin to swing in, thereby completing the nucleoside triphosphate binding pocket and activating the enzyme. Alternatively, the TTM and kinase domains may cooperatively bind a nucleoside triphosphate substrate and a second, yet to be described, metabolite to catalyze a novel phosphotransfer reaction. Mutation of conserved lysine and aspartate residues involved in UCK catalysis, inhibits the phosphotransferase activity of AtTTM1 *in vitro* and these residues are essential at least for AtTTM2 function *in vivo*, highlighting the functional significance of the enzymatic kinase module in AtTTM1/2. Structural analysis of the TTM domain itself revealed extensive structural similarities with bacterial/archaeal TTM enzymes. However, while the basic amino acids involved in binding of triphosphorylated substrates are conserved in AtTTM1/2, the invariant glutamates essential for the coordination of two metal co-factors are absent in this plant TTM-enzyme family. We speculate that the AtTTM1 and 2 β-tunnel domains may be able to specifically bind a tri- or polyphosphorylated metabolite. Although, previous studies have reported that both proteins act on pyrophosphate (18,19).

Our cellular studies demonstrate that the C-terminal transmembrane module tail-anchors both AtTTM1 and AtTTM2 to the outer mitochondrial membrane corroborating earlier studies (18,19). This localization distinguishes the plant proteins from other eukaryotic soluble TTM proteins. Very recently, AtTTM1 and 2 have been implicated in programmed cell death (20) during which the mitochondria play a critical role in providing energy but precise mechanisms of action have not yet been elaborated. Similarly the delay in bolting observed in *ttm2* mutants may point to mitochondrial involvement, as an alteration in energy metabolism is well documented during this process (27,28). On the other hand, AtTTM1 does not appear to be crucial for this developmental transition. These observations support the notion that these proteins have non-redundant biological roles (18) but the level at which this operates remains to be defined. Although both At*TTM1* and At*TTM2* are strongly expressed in shoot meristem tissue and are located to the same cellular organelle, it is interesting to note that the bolting delay in *ttm2* is rescued in transgenic plants expressing At*TTM2* under the constitutive and ubiquitous UBIQUITIN promoter (Fig. S5). We take this to indicate that a strict need for spatiotemporal tissue expression of At*TTM2* (i.e. in a specific place at a specific time) is not crucial for biological function. Thus, post-transcriptional events such as modification of the protein or interaction with a specific ligand/protein that facilitates pivoting of the helical hairpin to an active conformation, as deduced from the structure of TTM1, may ultimately account for functionality.

## MATERIALS AND METHODS

### Expression, purification and crystallization of AtTTM1

AtTTM1 and 2 constructs (residues 20-412) optimized for expression in *E. coli* (GeneArt, Thermofisher) were cloned into vector pMH-TrxT (providing an N-terminal octa-histidine-Streptavidin II-thioredoxin tag followed by a tobacco etch virus protease cleavage site). The recombinant plasmids were transformed in *E. coli* BL21 (DE3) RIL cells. Protein expression and purification are described in supporting material. Protein concentrations were determined using absorbance measurements at 280 nm using a NanoDrop™ OneC UV-Vis spectrophotometer (Thermo Fisher Scientific) and the estimated extinction coefficient (29). AtTTM1 crystals developed in hanging drops composed of 1 μL of protein solution (6 mg/ml AtTTM1 in 10 mM Na_2_HPO_4_/K_2_HPO_4_ pH 7.5, containing 250 mM NaCl, 0.5 mM Tris[2-carboxyethyl]phosphine) and 1 μl of crystallization buffer (30% PEG 8,000/20% ethylene glycol, 0.1 M carboxylic acids, 0.1 M Tris, pH 8.5, based on Morpheus condition G10 (30) suspended over 1 ml of the latter as reservoir solution. The adenosine-5’-[(β,γ)-methyleno]triphosphate (APPCP) complex was prepared by soaking crystals in crystallization buffer supplemented with 10 mM APPCP (Jena Biosciences), 10 mM uridine and 10 mM MgCl_2_ for 3 min. Crystals were directly snap frozen in liquid N_2_.

### Crystallographic data collection, structure solution and refinement

Redundant 3-wavelength multi-wavelength anomalous dispersion (MAD) data were collected at beamline PXIII of the Swiss Light Source, Villigen, Switzerland (Table S1) and processed and scaled with the programs XDS and XSCALE (31). Eight consistent Se sites were identified in SHELXD (32), site refinement and phasing was done in SHARP (33). A density modified electron density map generated with phenix.resolve (34) was readily interpretable and the model was built in alternating cycles of manual model building in Coot (35) and restrained refinement in phenix.refine against native data and a dataset recorded from a APPCP soaked crystal diffracting to 2.70 and 2.65 Å resolution, respectively (Table S1). Inspection of the final models with phenix.molprobity (36) revealed excellent stereochemistry (Table S1). Structural representations were done with Pymol (https://sourceforge.net/projects/pymol/) and ChimeraX (37).

### Plant material and growth conditions

*Arabidopsis thaliana* (Columbia ecotype), *ttm1-1* (SALK_079237, N579237), *ttm1-3* (SALK_126667, N626667), *ttm2-1* (SALK_145897, N645897), *ttm2-2* (SALK_114669, N614669), *ttm2-3* (SALK_084280, N584280) were obtained from the European Arabidopsis Stock Centre, whereas *ttm1-2* (SALK_GK-672E02) was obtained from the GABI-KAT collection (38). The *pTTM1:TTM1, pUBQ:YFP-TTM1, pUBQ:TTM1-YFP, pUBQ:YFP-TTM1*Δ*TM, pTTM1:GUS, pTTM2:TTM2, pUBQ:YFP-TTM2, pUBQ:TTM2-YFP, pUBQ:YFP-TTM2*Δ*TM* and *pTTM2:GUS* constructs were generated by amplifying the corresponding sequence of At*TTM1* or At*TTM2* (including 396 bp or 1551 bp upstream of the translational start codon in the case of *pTTM2:TTM1* and *pTTM2:TTM2*, respectively, or a truncated form of the coding sequence without the transmembrane domain in the case of *pUBQ:YFP-TTM1*Δ*TM* and *pUBQ:YFP-TTM2*Δ*TM* or the upstream region alone in the case of *pTTM1:GUS* and *pTTM2:GUS*) from Arabidopsis gDNA or cDNA of 10-day-old seedlings and cloned into the pENTR/D-TOPO vector using the pENTR TOPO cloning kit (Life Technologies) and subsequently cloned into Gateway® destination vector pBGW,0 (*pTTM1:TTM1*, p*TTM2:TTM2*) or pBGWFS7,0 (*pTTM1:GUS, pTTM2:GUS*) (39) or *pUBN-*YFP-DEST (40) by an LR reaction using LR clonase enzyme mix II (Life Technologies) to be expressed as AtTTM1 or AtTTM2, respectively, under the control of its own upstream promoter region (*pTTM1:TTM1, pTTM2:TTM2*), or as a fusion protein with YFP either at the N or C terminus under the control of the *ubiquitin10* promoter (At4g05320) (*pUBQ:YFP-TTM1, pUBQ:TTM1-YFP, pUBQ:YFP-TTM1*Δ*TM, pUBQ:YFP-TTM2, pUBQ:TTM2-YFP, pUBQ:YFP-TTM2*Δ*TM* respectively), or for expression of *GUS* under the control of the upstream region of *TTM1* (*pTTM1:GUS*) or *TTM2* (*pTTM2:GUS*). *pTTM2:TTM2* was used as a template to introduce point mutations in the P-loop kinase and TTM domains (annotated *pTTM2:TTM2 K78A D98A* and *pTTM2:TTM2 R299A R301A*, respectively) using the Quikchange Site-Directed Mutagenesis II XL Kit (Agilent Technologies) according to the manufacturers’ instructions.

### Tissue and cellular expression analysis

For histochemical localization of GUS activity several independent *pTTM1:GUS* and *pTTM2:GUS* lines were analyzed and treated as described in supporting material. Transient expression of *pTTM1:TTM1, pUBQ:YFP-TTM1, pUBQ:TTM1-YFP, pUBQ:YFP-TTM1*Δ*TM, pTTM2:TTM2, pUBQ:YFP-TTM2, pUBQ:TTM2-YFP, pUBQ:YFP-TTM2*Δ*TM* in Arabidopsis protoplasts was carried out as described in (41) using an SP5 confocal laser-scanning microscope (Leica Microsystems) equipped with a 63x oil NA 1.4 PlanApo objective. Fluorescence microscopy of tissues (cotyledon, hypocotyl, and root tissue) from stable lines carrying the same expression constructs was carried out on 4-day-old transgenic seedlings mounted in water between slide and coverslip with a double-sided scotch tape spacer (flow chamber). A 514 nm laser line was used to excite YFP and (when relevant) chlorophyll and to generate transmission images. YFP fluorescence was collected by a HyD detector between 519 nm and 560 nm, and chlorophyll fluorescence was collected by a PMT between 650 nm and 800 nm. When relevant, samples were incubated in 500 nM MitoTracker Red CMXRos (Invitrogen) for 30 min in water, then mounted in water in a flow chamber and excited at 579 nm for fluorescence collection by a HyD detector between 595 nm and 634 nm. Image analysis was performed using the software Fiji (42). A Gaussian blur of radius 0.6 pixels was applied to all z slices, and maximum intensity projections were applied and displayed.

## Data Availability

Crystallographic coordinates and structure factors have been deposited with the Protein Data Bank (http://rcsb.org) with accession codes 7Z66 (phosphate bound form) and 7Z67 (APPCP nucleotide analog bound form). The associated raw diffraction images have been deposited with zenodo.org (DOI:10.5281/zenodo.6365015, SeMAD datasets; DOI:10.5281/zenodo.6364933, native dataset for phosphate bound form; DOI:10.5281/zenodo.6365041, native dataset for APPCP nucleotide analog bound form). Other data generated for this study are included within this article and supporting information.

## Supporting Information

This article contains supporting information.

## ACKNOWLEDGMENTS

We gratefully acknowledge the European Research Council (ERC Starting Grant 310856, to M.H.), the Swiss National Science Foundation (grants 31003A-141117/1 and 31003A_162555/1 to T.B.F.), as well as the University of Geneva for supporting this work. We thank Mireille de Meyer-Fague for excellent dedicated support with Arabidopsis lines and Kitaik Lee for aspects of the protein purification; Luis López-Molina and Roman Ulm for kindly donating *ga1-3* and *co-101* single mutant lines, all University of Geneva.

