## Supporting Information for "Structural and functional insight into the plant unique multimodular triphosphosphate tunnel metalloenzymes of *Arabidopsis thaliana*"

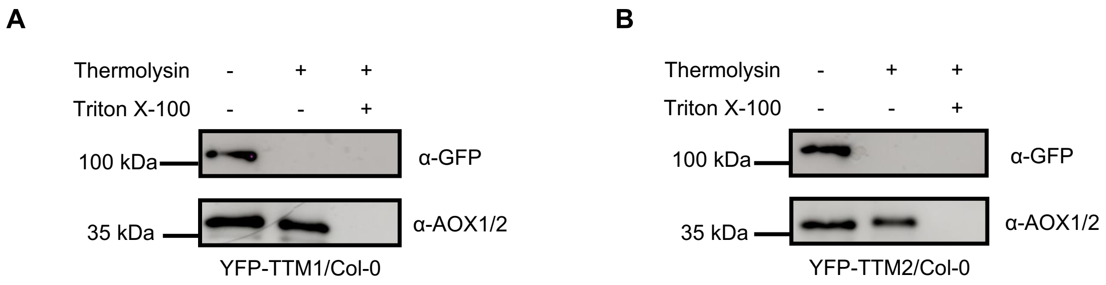


**Figure S1.** Immunoblot analyses of protease protection assays from purified intact mitochondria of Arabidopsis leaves expressing YFP-TTM1 (A) or YFP-TTM2 (B) untreated or treated with thermolysin (120 µmol/ml) in the absence or presence of 1% (v/v) Triton X-100 to disrupt mitochondrial membranes. YFP-TTM1 and YFP-TTM2 fusion proteins (102-kDa and 105-kDa, respectively) were detected with a GFP antibody (α-GFP). The intactness of mitochondria was assessed using an antibody against the inner mitochondrial membrane protein AOX (37-kDa) (α-AOX1/2). The antibody does not distinguish between the two isoforms AOX1 and AOX2. Numbers to the left of the panel indicate molecular mass in kilodaltons as a function of mobility.

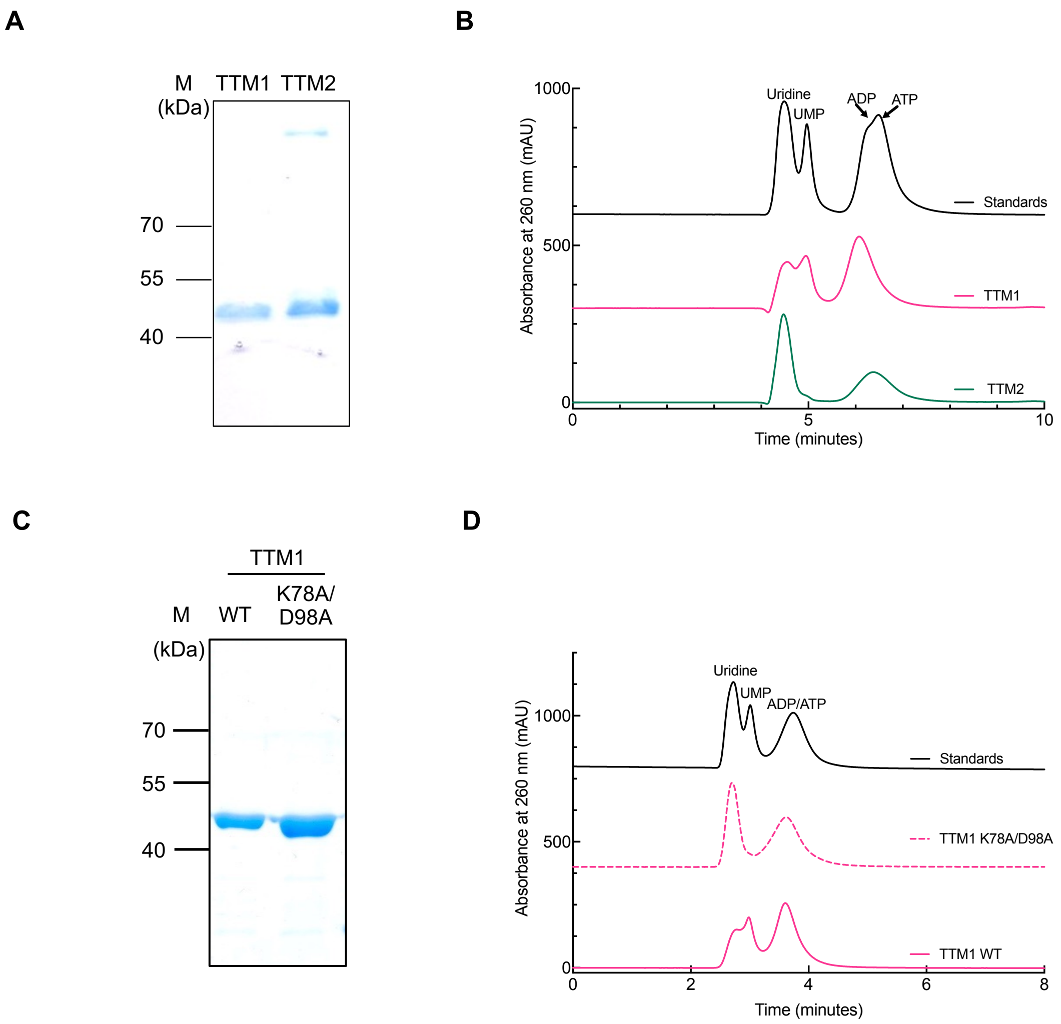


**Figure S2.** TTM uridine kinase activity. A, Purified tandem kinase and TTM modules (TTM1/$2^{20\text{-}412}$) of AtTTM1 or AtTTM2 (TTM1/TTM2, as indicated). B, HPLC analysis of phosphotransfer activity of TTM1 or TTM2 using uridine and ATP as substrates. Traces show the standards mixture (top); uridine, UMP, ATP or ADP (all 250 µM); or uridine, ATP (both 250 µM) in the presence of 2 µg of TTM1 (middle) or TTM2 (bottom). The TTM1 and standard profiles have been offset by 250 and 600 mAU to facilitate viewing. C, Purified tandem kinase and TTM modules ($TTM1^{20\text{-}412}$) wild type (WT) or K78A/D98A, as indicated. D, HPLC analysis of phosphotransfer activity as in B using 2 µg of TTM1 K78A/D98A (middle) or TTM1 WT (bottom). The TTM1 K78A/D98A and standard profiles have been offset by 400 and 800 mAU to facilitate viewing.

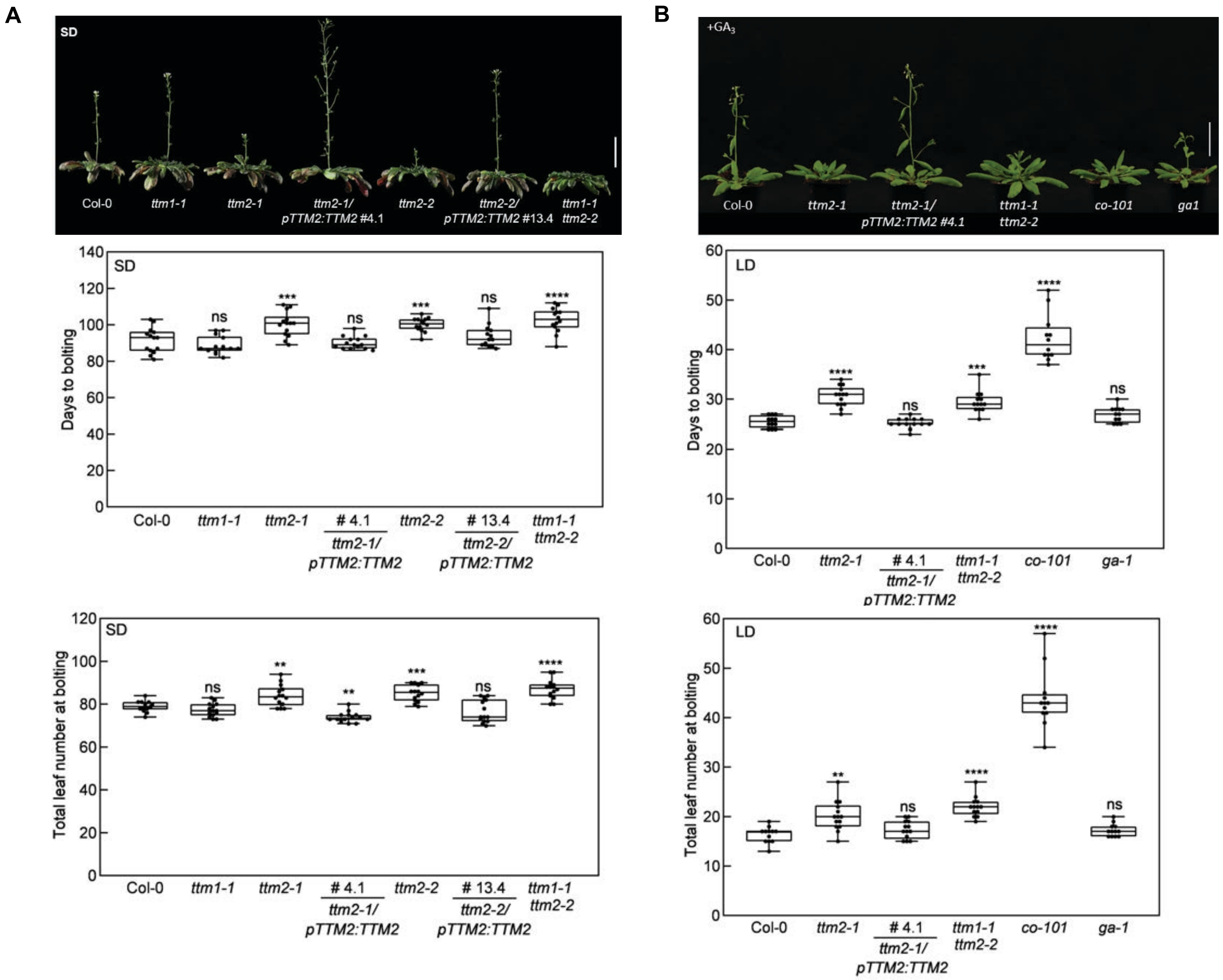


**Figure S3.** Analysis of the transition to the reproductive stage of *ttm1/2* alleles under an 8 hr photoperiod. A, Representative photographs of the transition to the reproductive stage of the lines as indicated compared to wild type (Col-0) under an 8 hr photoperiod (short day, SD), the scale bar = 5 cm, as well as the quantification of the switch to the reproductive stage represented by days to bolting and total leaf number (rosette and cauline leaves). Box-plot graphs show the median (horizontal bar in the box) and the minimum and maximum of the d ata (whisker extremities). One representative replicate of three biological replicates is shown (n = 12-14 per replicate). Statistical relevance was calculated by one-way ANOVA followed by Dunnett's multiple comparisons test, P > 0.05 ns, P<0.01**, P<0.001*** and P<0.0001****. B, Analysis of the effect of exogenous gibberellic acid (+$GA_3$) application on the reproductive transition. Representative photographs of wild type (Col-0), *ttm2*, *TTM2*-complemented (*pTTM2:TTM2*), *co-101* (CONSTANS mutant, delayed bolting control line), *ga-1* (GA biosynthesis mutant) grown in under a 16 hr photoperiod (LD) after treatment and quantification as in A.

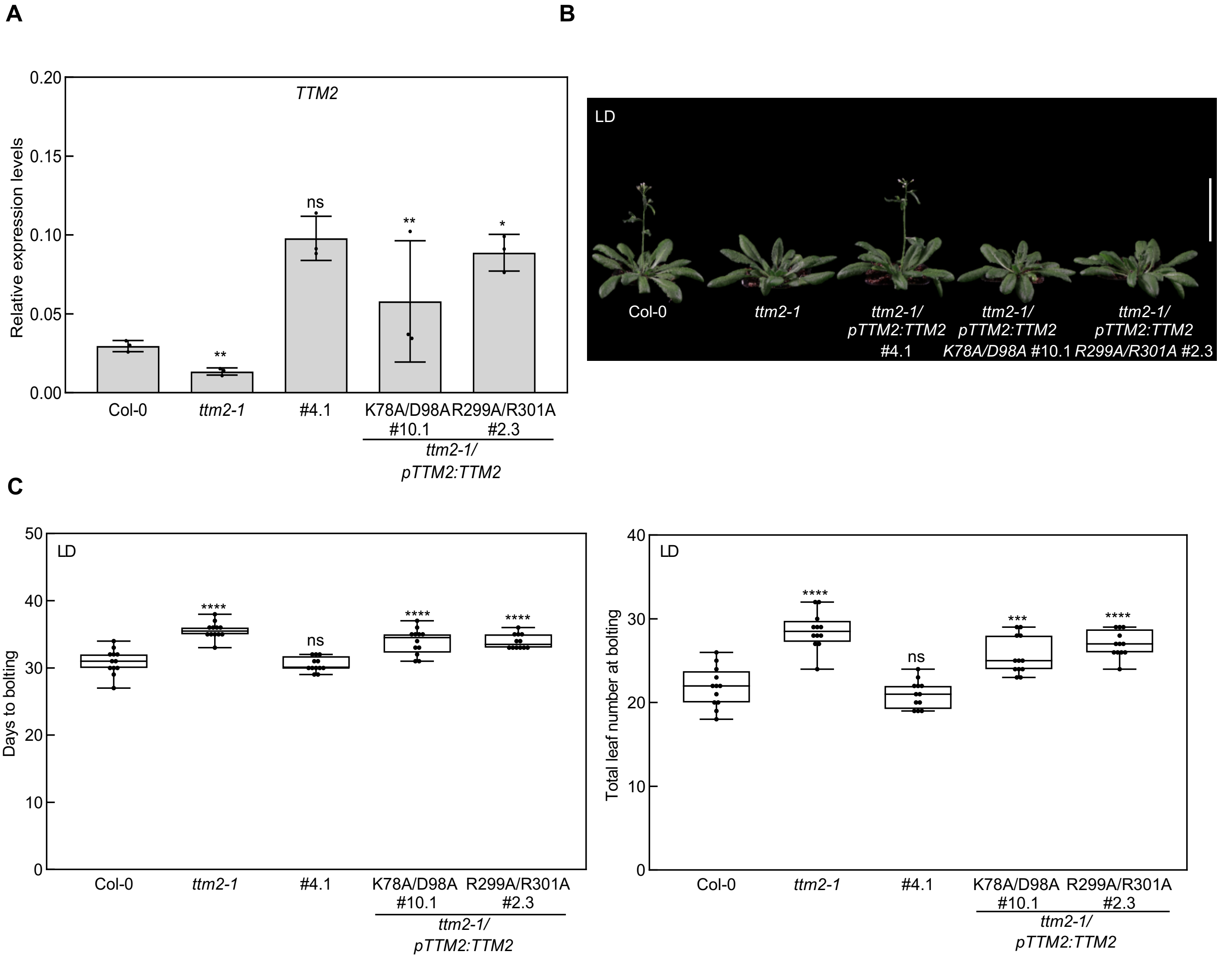


**Figure S4.** The kinase and TTM domains of AtTTM2 are required for functionality in bolting. A, Quantitative analysis of At*TTM2* expression in transgenic *ttm2-1* carrying the wild type (*pTTM2:TTM2*), kinase (*pTTM2:TTM2 K78A D98A*) or TTM (*pTTM2:TTM2 R299A R301A*) mutant alleles relative to *UBC21*. The data are the average of three independent biological replicates. Statistical relevance was calculated by a one-way ANOVA followed by Dunnett's multiple comparisons test; $P < 0.05$* and $P < 0.01$** and error bars represent standard deviation. B, Representative images of lines grown under a 16 hr photoperiod (LD). Scale bar = 5 cm. C, Quantification of the transition to the reproductive stage of *ttm2* lines as indicated compared to wild type (Col-0) grown under a 16 hr photoperiod (LD) represented by days to bolting and total leaf number (rosette and cauline leaves). Box-plot graphs show the median (horizontal bar in the box) and the minimum and maximum of the data (whisker extremities). One representative replicate of three biological replicates is shown (n = 12-14 per replicate). Statistical relevance was calculated by one-way ANOVA followed by Dunnett's multiple comparisons test, $P > 0.05$ ns, $P<0.001$*** and $P<0.0001$****. In all cases ns refers to no significance.

**A**

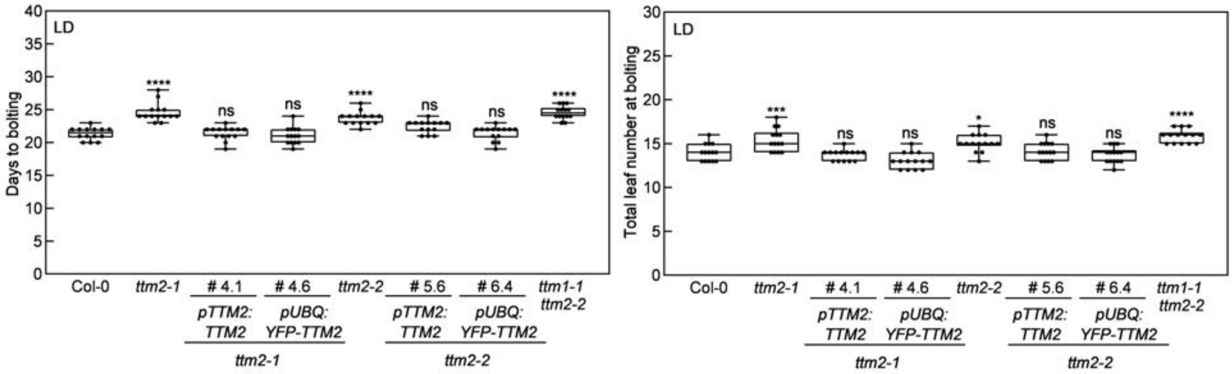


**B**

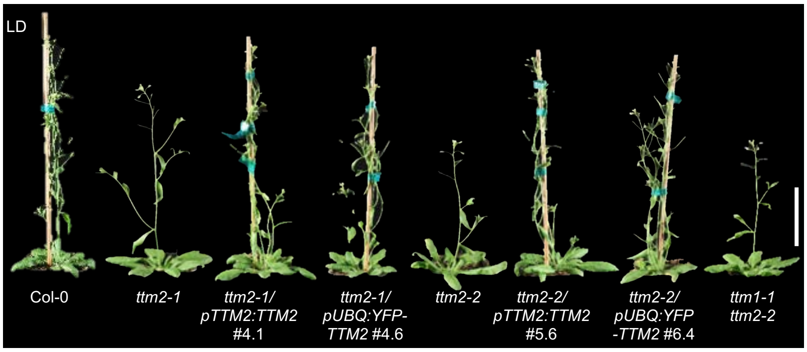


**Figure S5.** Expression of a translational fusion of YFP to the N-terminus of AtTTM2 under a constitutive and ubiquitous promoter is functional. A, Comparative quantification of the transition to reproductive stage of independent *ttm2* lines complemented with AtTTM2 alone under the control of its own upstream region (*pTTM2:TTM2*) or fused to YFP at the N-terminus under control of the UBIQUITIN promoter *(pUBQ:YFP-TTM2*), and *ttm1*/*ttm2* double mutant lines. Lines are compared to wild type (Col-0) grown under a 16 hr photoperiod (LD) represented by days to bolting and total leaf number (rosette and cauline leaves). Box-plot graphs show the median (horizontal bar in the box) and the minimum and maximum of the data (whisker extremities). One representative replicate of three biological replicates is shown (n = 12-14 per replicate). Statistical relevance was calculated by one-way ANOVA followed by Dunnett's multiple comparisons test, P > 0.05 ns, P<0.01**, P<0.001*** and P<0.0001****. Statistical relevance was calculated by one-way ANOVA followed by Dunnett's multiple comparisons test, P<0.05*, P<0.001*** and P<0.0001****. In all cases ns refers to no significance. B, Representative images of lines shown in A, scale bar = 5 cm.

**Table S1.** Data collection and refinement statistics for AtTTM1.

| | **AtTTM1 (inflection)** | **AtTTM1 (peak)** | **AtTTM1 (remote)** | **AtTTM1 (native)** | **AtTTM1 (APPCP)** |
|---|---|---|---|---|---|
| **PDB-ID** | | | | **7Z66** | **7Z67** |
| **Data collection** | | | | | |
| Energy (eV) | 12657.5 | 12665.0 | 12759.0 | | |
| Space group | | P $6_1$ 2 2 | | *P $6_1$ 2 2* | *P $6_1$ 2 2* |
| Cell dimensions | | | | | |
| *a*, *b*, *c* (Å) | | 93.83, 93.83, 266.79 | | 93.71, 93.71, 267.74 | 93.37, 93.37, 266.33 |
| α, β, γ (°) | | 90,90,120 | | 90, 90, 120 | 90, 90, 120 |
| Resolution (Å) | | 49.88 – 2.91 (3.08 – 2.91) | | 46.86 – 2.70 (2.87 – 2.70) | 45.69 – 2.65 (2.81 – 2.65) |
| $R_{meas}$[#] | 0.389 (4.86) | 0.368 (4.22) | 0.42 (5.6) | 0.157 (2.51) | 0.163 (2.80) |
| CC(1/2)[#] | 0.99 (0.24) | 1.0 (0.30) | 1.0 (0.18) | 1.0 (0.58) | 1.0 (0.65) |
| $I/\sigma I$[#] | 10.7 (0.8) | 11.2 (0.8) | 9.9 (0.6) | 14.3 (1.2) | 17.5 (1.48) |
| Completeness (%)[#] | 99.9 (99.2) | 99.8 (98.9) | 99.8 (99.1) | 99.9 (99.3) | 99.7 (99.1) |
| Redundancy[#] | 21.4 (21.0) | 21.4 (21.0) | 21.4 (21.2) | 21.2 (21.2) | 19.2 (18.3) |
| Wilson B-factor[#] | 76.0 | 74.7 | 77.5 | 83.2 | 82.5 |
| **Refinement** | | | | | |
| Resolution (Å) | | | | 48.86 – 2.70 | 45.69 – 2.65 |
| No. reflections | | | | 35,942 | 37,731 |
| $R_{work}/R_{free}$[$] | | | | 0.212 (0.238) | 0.219 (0.245) |
| No. atoms | | | | | |
| protein | | | | 3,169 | 3,174 |
| buffer | | | | 22 | 50 |
| solvent | | | | 29 | 4 |
| Res. B-factors[$] | | | | | |
| protein | | | | 96.5 | 92.7 |
| buffer/APPCP | | | | 97.2 | 109.2 |
| solvent | | | | 79.8 | 69.5 |
| R.m.s deviations[$] | | | | | |
| bond lengths (Å) | | | | 0.0023 | 0.0023 |
| bond angles (°) | | | | 0.48 | 0.49 |
| Ramachandran plot[$]: | | | | | |
| most favored regions (%) | | | | 98.6 | 98.7 |
| outliers (%) | | | | 0 | 0 |
| MolProbity score[$] | | | | 1.68 | 1.68 |

[#]as defined in XDS ( https://doi.org/10.1107/S0021889893005588)
[$]as defined in phenix.refine/phenix.molprobity (https://doi.org/10.1093/nar/gkm216)

**Table S2.** List of primers used in this study.

| | Purpose | Gene locus | Oligonucleotide sequence (5′→ 3′) | |
|---|---|---|---|---|
| | | | Forward primer | Reverse primer |
| Genotyping | *ttm2-1* | At1g26190 | CTTCTGGTGCTGGAAAGACT | TATGTCCATCAGAAAGGACC |
| | *ttm2-2* | | GTATCCGTAACCCATTCCTGG | GTAATGCGTGACGTGATTGT |
| | *ttm2-3* | | TGAAGGATCAAGTCCAATTG | CAAGATGTAAGTCGGGCTCT |
| | *ttm1-1* | At1g73980 | TTCTGCTCATGCTTTGATTTG | AGCAAGGGTGATTAATCTGG |
| | *ttm1-2* | | CTTCCATCAGAGGAGATTCC | CTTGGAGGATTAATGGCACT |
| | *ttm1-3* | | AAAATGTGGCAGACCAACTT | ATGCATTATCTGACGCTGTCC |
| | *tom20-2* | At1g27390 | AATCCAAGGACTTGCGATAT | TTTGTTCATATGGGCCTTCTG |
| T-DNA | SALK | | GCGTGGACCGCTTGCTGCAA | |
| | GABI-KAT | | ATATTGACCATCATACTCATT | |
| Cloning | *pTTM2*:GUS | At1g26190 | *CACC*GAAAGGCAGAAAGGTAGTCG[a] | GACCCATTATGAACAATAATCT |
| | *pTTM2*:TTM2 | | *CACC*GAAAGGCAGAAAGGTAGTCG[a] | AAACTAAAAACCTAAACTTTATTGAATA |
| | YFP-TTM2 | | *CACC*ATGGGTCAAGACAGCAATGG[a] | TTATTGCCGCTTGTTAATGTAG |
| | TTM2-YFP | | *CACC*ATGGGTCAAGACAGCAATGG[a] | TTGCCGCTTGTTAATGTAGCTC |
| | YFP-TTM2ΔTM | | *CACC*ATGGGTCAAGACAGCAATGG[a] | CTACTTGTTAGATTCCACATTGG |
| | *pTTM1*:GUS | At1g73980 | *CACC*ACCCCAATAAGGATTCTACTCG[a] | GCCTCAAAGTTGCAAATGAAG |
| | YFP-TTM1 | | *CACC*ATGGCGCTCGATAGTTCTG[a] | CTATTTCAGACGACTGTAGGCA |
| | TTM1-YFP | | *CACC*ATGGCGCTCGATAGTTCTG[a] | TTTCAGACGACTGTAGGCAAA |
| | YFP-TTM1ΔTM | | *CACCC*ATGGCGCTCGATAGTTCTG[a] | CTAAGTAGAGGATTCGAGGTTTT |
| Mutagenesis | *gDNA* K78A | At1g26190 | CCTTCTGGTGCTGGAGCAACTGTAT<br>TCACTGAG | CTCAGTGAATACAGTTGCTCCAGCACC<br>AGAAGG |
| | *gDNA* D98A | | GCTGTCATTTCAATGGCTAATTATA<br>ATGATTCTAGTCGG | CCGACTAGAATCATTATAATTAGCCAT<br>TGAAATGACAGC |
| | *gDNA* R299A | | CAGAGTCGTGCCAATCATATTTGGC<br>GATGCGGAATAAAGATGG | CCATCTTTATTCCGCATCGCCAAATATG<br>ATTGGCACGACTCT |
| | *gDNA* R301A | | GTGCCAATCATATTTGAGGATGGCG<br>AATAAAGATGGAAAGTACAGC | GCTGTACTTTCCATCTTTATTCGCCATC<br>CTCAAATATGATTGGCA |
| | TTM2 K78A | | TTTCGGTAAACACGGTTGCACCTGC<br>ACCGCTCGGAC | GTCCGAGCGGTGCAGGTGCAACCGTG<br>TTTACCGAAA |
| | TTM2 D98A | | GGCTGCTATCATTATAGTTAGCCAT<br>GCTAATAACGGCAACG | CGTTGCCGTTATTAGCATGGCTAACTA<br>TAATGATAGCAGCC |
| | TTM2 R299A | | CGGAAAGCTGTCAGAGCTATCTGGC<br>TATGGCTAATAAAGATGGTAAATAT<br>TCCC | GGGAATATTTACCATCTTTATTAGCCA<br>TAGCCAGATAGCTCTGACAGCTTTCCG |
| | TTM2 R301A | | CTGTCAGAGCTATCTGCGTATGGCT<br>AATAAAGATGGTAAATATTCCC | GGGAATATTTACCATCTTTATTAGCCA<br>TACGCAGATAGCTCTGACAG |
| | TTM1 K78A | At1g73980 | AGGTCCGAGCGGTGCAGGTGCAAC<br>CATTTTTACCGAAAAAA | TTTTTTCGGTAAAAATGGTTGCACCTG<br>CACCGCTCGGACCT |
| | TTM1 R299A | | GCATGTCAGAGCTATCTGGCTATGG<br>CTAATCGTGATGG | CCATCACGATTAGCCATAGCCAGATA<br>GCTCTGACATGC |
| | TTM1 R301A | | GTCAGAGCTATCTGCGTATGGCTAA<br>TCGTGATGGCAAATATA | TATATTTGCCATCACGATTAGCCATAC<br>GCAGATAGCTCTGAC |

| | | | Forward primer | Reverse primer |
|---|---|---|---|---|
| qPCR | *4F/R* | At1g26190 | TGAAGATCCAGAGTCGTGCCAAT | AGGAGTATCCGTAACCCATTCCT |
| | *2F/R* | At1g73980 | TTATCATTCTCCGCATCCGC | GCAACCAGTACGACCTCTTC |
| | *UBC21* | At5g25760 | TAGCATTGATGGCTCATCCTG | TTGTGCCATTGAATTGAACCC |
| | *PP2AA3* | At1g13320 | TAACGTGGCCAAAATGATGC | GTTCTCCACAACCGCTTGGT |
| RT | *TTM2* | At1g26190 | | TCGAGAACGTCTTCCAAACG |
| | *TTM1* | At1g73980 | | TGCCACTATTTCACGAAGCC |
| | *PP2AA3* | At1g13320 | | GCACCAAAAAGCAAATACGC |

[a] The four nucleotides added for directional cloning in pENTR/D-TOPO vector are depicted in italics

**Recombinant *TTM1/2* expression, purification and HPLC assay.**

For AtTTM1 protein production in *E. coli* BL21 (DE3) RIL cells (Invitrogen) for crystallization, an overnight culture in lysogeny broth (LB) growth medium was pelleted by centrifugation (4,500 *g*) and resuspended in 1 liter minimal M9 medium consisting of 20 mM $NH_4Cl$, 8.5 mM NaCl, 47 mM $Na_2HPO_4$, 22 mM $KH_2PO_4$, 30 µM $FeSO_4$, 1 mM $MgSO_4$, 0.1 mM $CaCl_2$, 0.4% (w/v) glucose and 1 mg each of thiamine, riboflavin, niacinamide and pyridoxine. The cells were grown at 37°C to $OD_{600}$=0.4 and methionine biosynthesis was inhibited by the addition of 50 mg leucine, 50 mg isoleucine, 50 mg valine, 100 mg phenylalanine, 100 mg lysine and 100 mg threonine. After 30 min, 60 mg of L-selenomethionine and 0.5 mM isopropyl β-D-1-thiogalactopyranoside (IPTG) were added to induce expression of selenomethionine labeled AtTTM1, and cells were grown for 3 hr at 37°C and for an additional 8 hr at 18°C. Cell pellets were collected by centrifugation at 4,500 *g* for 30 min, resuspended in lysis buffer (50 mM sodium phosphate, pH 7.5, containing 500 mM NaCl, 5 mM β-mercaptoethanol*,* 1 mM phenylmethylsulfonyl fluoride, and Complete Protease Inhibitor Cocktail Roche), and disrupted by sonication. The cell suspension was spun down at 18,000 *g* for 1 hr at 4°C, and the supernatant was sequentially loaded onto Ni-NTA (HisTrap HP 5 ml; GE Healthcare) and Strep (Strep-Tactin XT Superflow; IBA Lifesciences) affinity columns. Proteins were eluted in buffer A (50 mM sodium phosphate pH 7.5, containing 500 mM NaCl) supplemented with 500 mM imidazole and 1x Buffer BXT (Strep-Tactin®XT elution buffer containing 50 mM biotin (IBA Lifesciences), respectively. The thioredoxin affinity tag was cleaved overnight with tobacco etch virus (TEV) protease at 4°C. An additional Ni-NTA affinity chromatography step was performed to remove the TEV protease and the cleaved affinity tag. Proteins were purified to homogeneity by size-exclusion chromatography on a HiLoad 26/600 Superdex 200 pg column (GE Healthcare) in buffer (25 mM Tris-HCl pH 8.0, containing 50 mM NaCl and 10 mM $MgCl_2$). For biochemical assays, heterologous protein expression was carried out in LB growth medium using 0.1 mM IPTG and incubation at 18˚C for 14-18 hr. AtTTM1 was purified as above by sequential Ni-NTA and Strep affinity chromatography. Point mutations of the kinase (K78A and D98A) were carried out using the Quikchange Site-Directed Mutagenesis II XL Kit (Agilent Technologies) according to the manufacturers' instructions and confirmed by sequencing (Microsynth). Primers used are listed in Table S2. Protein expression and purification was as described above. AtTTM2 was purified by Ni-NTA (Protino Ni-NTA, Macherey-Nagel; equilibrated in 50 mM sodium phosphate, pH 7.5, containing 300 mM NaCl and 10 mM imidazole; supplemented with 300 mM imidazole for elution). After removal of imidazole by dialysis in 50 mM sodium phosphate, pH 7.5, containing 300 mM NaCl, TEV cleavage of the tag was carried out in the same buffer at 4˚C for 14 hr followed by repetition of the Ni-NTA chromatography step. The proteins were further purified by size-exclusion chromatography (Superdex 200 increase 10/300 GL, GE Healthcare; equilibrated in 50 mM sodium phosphate, pH 7.5, containing 200 mM NaCl) before use. Phosphotransfer activity was carried out with 2 µg of purified protein in 25 mM Tris-HCl, pH 8.0, containing 250 mM NaCl, 10 mM $MgCl_2$, 10 mM KCl and 250 µM each of uridine and ATP in a total volume of 30 µl at 25°C for 30 minutes. Protein was precipitated by heating and centrifugation and the supernatant was used for analysis by HPLC equipped with a diode array detector. HPLC analysis was carried out on a VYDAC® reversed-phase HPLC column (#201HS52, Thermo Fisher Scientific) using an injection volume of 20 µl and elution with an isocratic gradient of 100 mM potassium phosphate pH 6.5, containing 10 mM tetrabutyl ammonium bromide and 5% acetonitrile at a flow rate of 0.25 mL per minute. Compounds were detected based on their absorbance at 260 nm.

**Plant material and growth conditions**

The *ga1-3* and *co-101* single mutant lines were gifts from Luis López-Molina and Roman Ulm, respectively (University of Geneva, Switzerland). *ttm1-1* and *ttm2-2* were crossed to generate *ttm1-1 ttm2-2*. In all cases the Columbia ecotype was used as wild type and all mutant lines were isolated to be in the homozygous state based on the corresponding antibiotic resistance, as well as PCR genotyping and sequencing. The primers used are listed in Table S2. In general, plants were grown on soil or sterile culture under 16 hr or 8 hr photoperiods as specified (60 % relative humidity, 120-150 µmol photons $m^{-1}$ $s^{-1}$ generated by fluorescent lamps (Philips Master T-D Super 80 18W/180 and 22°C) followed by the corresponding hours of darkness (8 or 16 hr) at 18°C and ambient $CO_2$ to complete a diel cycle. Seeds were stratified for 2-3 days in the dark at 4°C before transferring to a growth incubator. Seeds used for *in vitro* culture were surface sterilized and air-dried prior to plating on half-strength Murashige and Skoog medium (Duchefa, M0221) in 0.55% [w/v] agar (Duchefa, P1001) plates (½ MS agar plates) (1). Seeds cultivated on soil (Einheitserde, Classic Ton Kokos) were grown in a CLF Climatics AR-66 phytochamber, whereas a CLF Climatics CU-22 L phytochamber was used for sterile cultures.

**Mitochondria isolation and thermolysin treatment.**

Mitochondria were isolated according to (2) with minor modifications. Around 100 g of Arabidopsis leaf material of plants grown under a 16 hr photoperiod was homogenized with grinding buffer (25 mM $Na_4P_2O_7$, 10 mM $KH_2PO_4$, pH 7.5 containing 0.3 M sucrose, 2 mM EDTA, 1% [w/v] BSA, 1% [w/v] PVP-40 and 20 mM ascorbic acid). A Waring blender was used 3-4 x 5 s at low speed with a ratio of at least 4 ml grinding buffer/g plant material. The suspension was filtered through four layers of muslin (Miracloth, Calbiochem®) via a funnel into a cooled beaker. The filtered homogenate was poured into 50-ml falcon tubes and centrifuged at 2,500 *g* for 5 min in a Sorvall Legend X1R centrifuge at 4°C. The supernatant was transferred to new tubes and spun down at 17,500 *g* for 18 min at 4°C. The supernatant was discarded and the pellet was resuspended in 10 ml of wash medium

(10 mM MOPS-KOH, pH 7.2 containing 0.3 M sucrose, 1 mM EDTA and 1% [w/v] BSA) with the aid of a soft bristle paint brush. The resuspended organelles were transferred to 50 ml falcon tubes. The volume was adjusted with wash medium to a final volume of 40 ml and samples were centrifuged at 3,200 *g* for 5 min. The supernatant was transferred into new tubes and the organelles were sedimented by centrifugation at 17,500 *g* for 18 min. The resulting supernatant was discarded and the washed organelles were resuspended uniformly in ~1 ml of wash medium, as above. Washed mitochondria were layered over 23 ml of a 28% [v/v] continuous gradient of Percoll® (Sigma) in wash medium in a 25 ml tube and centrifuged at 26,200 *g* for 50 min in an Oprima XPN80 ultracentrifuge (Beckman Coulter). Mitochondria were aspirated with a Pasteur pipette avoiding collection of the yellow or green plastid fractions. The suspension was diluted with 4 volumes of standard wash medium and centrifuged at 17,500 *g* for 18 min in 35 ml centrifuge tubes. The resultant loose pellet was resuspended in wash medium and centrifuged again at 17,500 *g* for 18 min. Finally, the mitochondrial pellet was resuspended in wash medium and the protein concentration was estimated using a NanoDrop™ OneC UV-Vis spectrophotometer (Thermo Fisher Scientific). Thermolysin treatment was performed as described in (3). Briefly, 40 µl of mitochondria purified from a *pUBQ:TTM1-YFP* or *pUBQ:TTM2-YFP* line were washed with 200 µl wash buffer (50 mM HEPES pH 7.5 containing 0.33 M sorbitol) and centrifuged at 16,000 *g* for 2 min at 4°C. The pellet was resuspended in 100 µl wash buffer and 120 $\mu g.ml^{-1}$ thermolysin (1 $mg.ml^{-1}$ stock solution) from *Bacillus thermoproteolyticus* (Fluka) with either 0.5 mM calcium chloride or water (as a negative control). The reaction was performed at 4°C for 30 min and stopped by adding 10 mM EDTA and incubation for 5 min. The reaction was centrifuged at top speed in a table top centrifuge (Eppendorf) and resuspended in 50 µl of SDS-PAGE buffer (63 mM Tris-HCl, pH 6.8 containing 0.1% [v/v] β-mercaptoethanol, 0.0005% [w/v] Bromophenol blue, 10% [v/v] glycerol and 2% [v/v] SDS).

**Tissue expression analysis**

For histochemical localization of GUS activity several independent *pTTM1:GUS* and *pTTM2:GUS* lines were analyzed and treated as follows: seedlings or tissues were collected in Eppendorf tubes filled with ice-cold 90% acetone and incubated for 20 min at room temperature; the plant material was washed three times with staining buffer (10 mM $NaH_2PO_4$/$Na_2HPO_4$ buffer pH 7.0, containing 0.5 mM $K_3[Fe(CN)_6]$, 0.5 mM $K_4[Fe(CN)_6]$, 0.1% [v/v] Triton X-100) and afterwards infiltrated for 12-16 hr at 37ºC in the dark with 5-bromo-4-chloro-3-indolyl-β-D-glucuronic acid (Sigma) in staining buffer (0.1 $mg.ml^{-1}$). Excess staining solution and chlorophyll was removed from the samples by rinsing with ethanol in the series 20%, 35% and 50% [v/v] for 30 min each time. Samples were then fixed in a solution containing 5% [v/v] formaldehyde, 10% [v/v] acetic acid, 50% [v/v] ethanol for

30 min and soaked in 70% [v/v] ethanol as a final dehydration step. Images were captured using a Leica MZ16 stereomicroscope equipped with an Infinity 2 digital camera (Leica Microsystems).

**Immunochemical analyses.**

Forty µl of samples from isolated mitochondria were boiled for 3 min in SDS-PAGE buffer (63 mM Tris-HCl, pH 6.8 containing 0.1% [v/v] β-mercaptoethanol, 0.0005% [w/v] Bromophenol blue, 10% [v/v] glycerol and 2% [v/v] SDS). Proteins were separated by SDS-PAGE (10%) before being transferred to nitrocellulose membranes using the iBlot system (Invitrogen) applying 20 volts for 7 min. The membranes were blocked for 1 hr in 10 mM Tris-HCl pH 7.4 containing 150 mM NaCl, 0.1% [v/v] Tween-20 (TBS-T) and 5% [w/v] milk powder (TBS-T milk solution) followed by incubation overnight at 4°C with the indicated primary antibody in TBS-T milk solution. After three wash steps for 10 min each using TBS-T, the membrane was incubated with secondary peroxidase-conjugated goat anti-mouse or anti-rabbit antibody, as appropriate (Bio-Rad) in TBS-T milk solution for 1 hr and washed three times for 10 min in TBS-T. The primary and secondary antibodies were used at dilutions of 1:1000 and 1:5000, respectively. Detection was carried out by chemiluminescence using Western Bright ECL (Advansta) and images were captured using an ImageQuant LAS 4000 system (GE Healthcare). The primary antibody against AOX1/2 was from Agrisera, and that against GFP from Santa-Cruz Biotechnology.

**Gene expression analysis by qPCR.**

Total RNA was extracted from 100-150 mg of frozen ground shoot material of 14-day old seedlings maintained on culture plates under a 16 hr photoperiod using the RNA NucleoSpin Plant kit (Macherey-Nagel) following the protocol provided by the company and treated with RNase-free DNase I to remove traces of DNA. A total of 1 µg of RNA was treated a second time with DNase RQ1 (Promega) and reverse transcribed into cDNA using Superscript II reverse transcriptase (Life Technologies) and either oligo(dT)$_{15-18}$ primers (Promega) or specific RT primers according to the manufacturer's recommendations. Quantification analyses were performed in 384-well plates using a QuantStudio5 Fast real-time PCR instrument (Applied Biosystems) by fluorescence-based real-time PCR using Power(Up) SYBR Green master mix (Applied Biosystems) and the following amplification program: 10 min denaturation at 95°C followed by 40 cycles of 95°C for 15 s and 60°C for 1 min. The data were analyzed using the comparative cycle threshold method ($2^{-\Delta CT}$) normalized to the reference gene *UBC21 (*At5g25760) or *PP2AA3* (At1g13320). Primers used are listed in Table S2. Each experiment was performed with at least three biological and three technical replicates.

**Giberellin treatment.**

The bioactive gibberellic acid $GA_3$ (Sigma) was prepared in ethanol (20 mM). The Arabidopsis mutant line *ga1-3* needs exogenous gibberellins to germinate and was used as a positive control. Seeds were sterilized with ethanol and incubated with 10 µM $GA_3$ in water solution-moistened filter paper for 2 days in darkness. After stratification, seeds were rinsed with water before sowing them on soil. Exogenous $GA_3$ during vegetative growth under a 16 hr photoperiod was applied twice weekly by spraying plants with a solution of 50 µM $GA_3$ in water and 0.02% [v/v] Tween-20**.**

**Statistical Analyses.**

Data was analyzed using one way-ANOVA with post-hoc Dunnet's multiple comparison test using GraphPad Prism software. The means and standard deviations were derived from replicated independent biological samples. Asterisks indicate statistically significant differences as annotated.
